# A tutorial on how (not) to over-interpret STRUCTURE/ADMIXTURE bar plots

**DOI:** 10.1101/066431

**Authors:** Daniel J Lawson, Lucy van Dorp, Daniel Falush

**Affiliations:** University of Bristol, Integrative Epidemiology Unit, Population Health Sciences, Bristol, UK; University College London Genetics Institute (UGI). Dept. Genetics, Evolution and Environment. London, UK; Centre for Mathematics and Physics in the Life Sciences and Experimental Biology (CoMPLEX), University College London. London, UK; Milner Centre for Evolution, University of Bath

## Abstract

Genetic clustering algorithms, implemented in popular programs such as STRUCTURE and ADMIXTURE, have been used extensively in the characterisation of individuals and populations based on genetic data. A successful example is the reconstruction of the genetic history of African Americans who are a product of recent admixture between highly differentiated populations. Histories can also be reconstructed using the same procedure for groups which do not have admixture in their recent history, where recent genetic drift is strong or that deviate in other ways from the underlying inference model. Unfortunately, such histories can be misleading. We have implemented an approach (badMIXTURE, available at github.com/danjlawson/badMIXTURE) to assess the goodness of fit of the model using the ancestry “palettes” estimated by CHROMOPAINTER and apply it to both simulated data and real case studies. Combining these complementary analyses with additional methods that are designed to test specific hypotheses allows a richer and more robust analysis of recent demographic history based on genetic data.

## INTRODUCTION

### STRUCTURE/ADMIXTURE are excellent tools for analysing recent admixture between differentiated groups

Model-based clustering has become a popular approach to visualize the genetic ancestry of humans and other organisms. Pritchard et al. [1] introduced a Bayesian algorithm STRUCTURE for defining populations and assigning individuals to them. FRAPPE and ADMIXTURE were later implemented based on a similar underlying inference model but with algorithmic refinements that allow them to be run on datasets with hundreds of thousands of genetic markers [2, 3].

One motivating example for the algorithms was African Americans. The “admixture model” of STRUCTURE assumes that each individual has ancestry from one or more of *K* genetically distinct sources. In the case of African Americans, the most important sources are West Africans, who were brought to the Americas as slaves, and European settlers. The two groups are thought to have been previously separated with minimal genetic contact for tens of thousands of years. This means that their history can be separated into two phases, a “divergence phase” lasting thousands of years of largely independent evolution and an “admixture phase”, in which large populations met and admixed within the last few hundred years. Specifically, most of the ancestors of African Americans that lived 500 years ago were either Africans or Europeans. The goal of the algorithm is to reconstruct the gene frequencies of these two distinct “ancestral” populations and to estimate what proportion of their genome each African American inherited from them (see Case study 1).

#### Case study 1: African Americans

When the STRUCTURE admixture model is applied to a dataset consisting of genetic markers from West Africans, African Americans and Europeans it infers two ancestral populations [1]. Each of the Europeans and Africans are assigned a great majority of their ancestry from one of them. African-Americans are inferred to have an average of 18% ancestry from the European cluster but with substantial inter-individual variation [4].

Assignment of clusters in this case is readily biologically interpretable. There are of course genetic differences amongst both the Africans and the Europeans who contributed to African American ancestry, e.g. reflecting genetic variation between regions within Europe and Africa, but the divergence between Europeans and Africans took place over millennia and is of a different magnitude to the recent admixture. These subtle differences are for example likely to have a relatively minor effect on the amount of African and European ancestry estimated for each individual, making an interpretation of the STRUCTURE admixture proportion as an estimate of the recent admixture fraction a reasonably accurate one.

### Qualitatively different historical scenarios can give indistinguishable STRUCTURE/ADMIXTURE plots

The STRUCTURE barplot has become a de-facto standard used as a non-parametric description of genetic data [5] alongside a Principle Components Analysis [6]. Case Study 1 and other successful examples of inference [7–9], as well as the difficulty of interpreting the results at all if they are not taken literally, have led most researchers to be cautious but literal with their interpretation, as caricatured in Figure 1 and described in case studies to follow. Many real population histories are not neatly separable into divergence and admixture phases but the methods can be applied to all datasets to produce ancestry bar plots.

**Figure 1:**
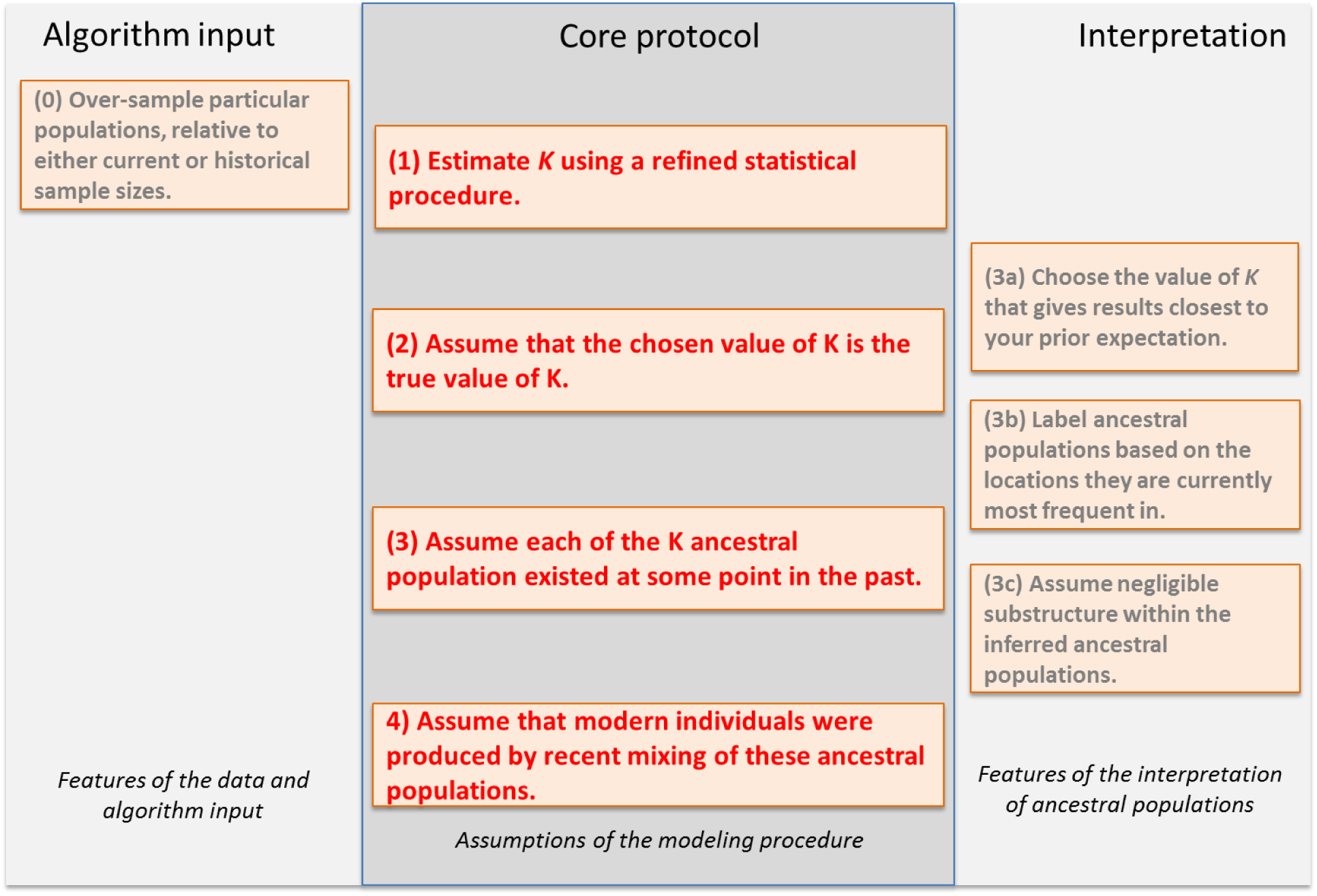
A protocol for interpreting admixture estimates, based on assuming that the model underlying the inference is correct. If these assumptions are not validated, there is a substantial danger of over-interpretation. The “Core protocol” describes the assumptions that are made by the admixture model itself (Protocol 1,3,4), and inference for estimating K (Protocol 2). The “Algorithm input” protocol describes choices that can further bias results, while the “Interpretation” protocol describes assumptions that can be made in interpreting the output that are not directly supported by model inference.

Figure 2 shows admixture histories inferred by STRUCTURE for three demographic scenarios. These simulations were performed with 13 populations (see Methods) – which provides valuable out-group information - but only results for the four most relevant populations are shown. The “Recent Admixture” scenario represents a history qualitatively similar to African Americans, in which the admixture model holds. The true history is that P2 is an admixture of P1, P3 and P4. ADMIXTURE, interpreted according to the protocol, infers that this is what happened and estimates approximately correct admixture proportions (true admixture proportions are 35% light green and 15% light pink).

**Figure 2.**
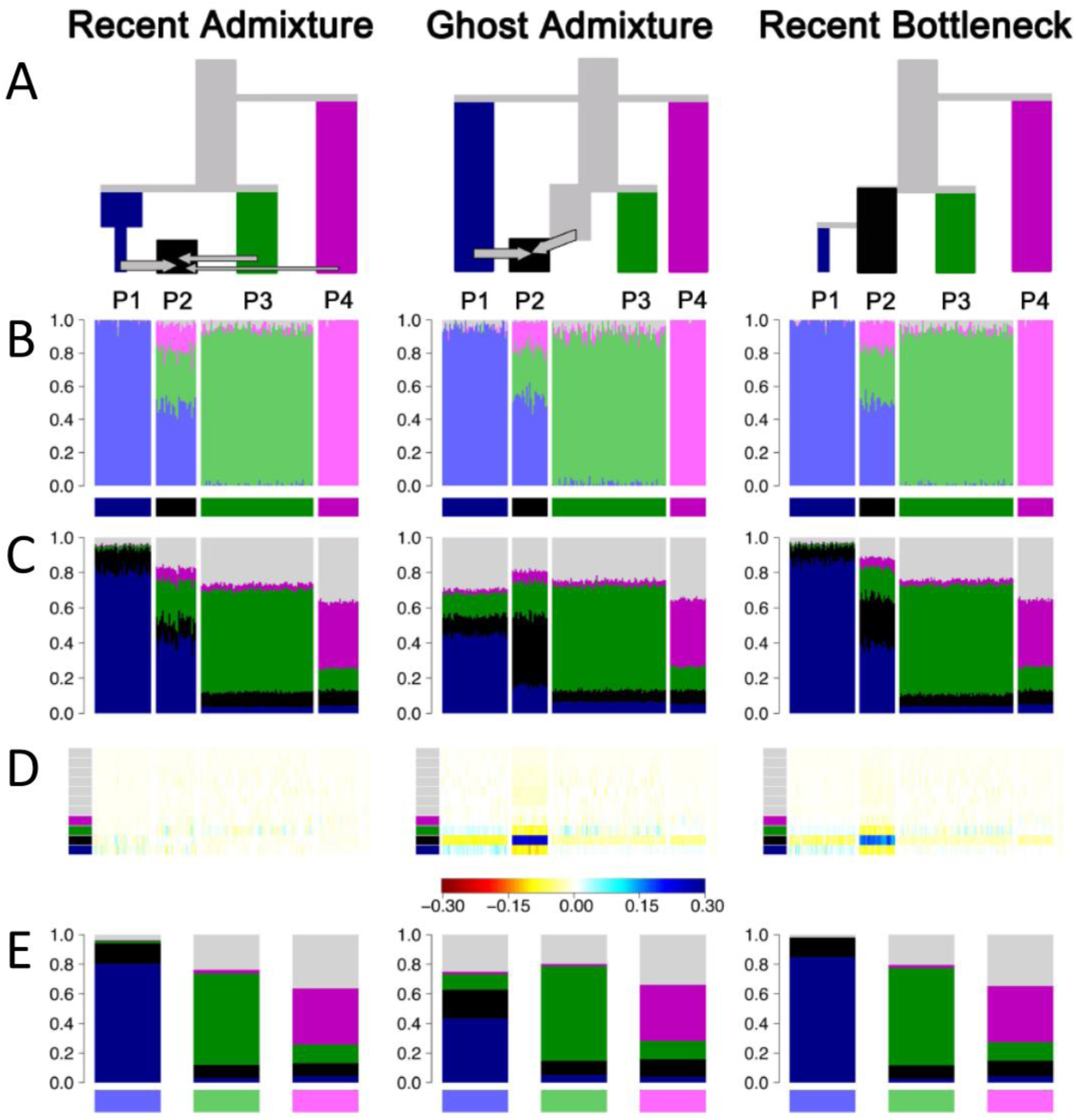
Three scenarios that give indistinguishable ADMIXTURE results. (A) Simplified schematic of each simulation scenario. (B) Inferred ADMIXTURE plots at K=11. (C) CHROMOPAINTER inferred painting palettes. (D) Painting residuals after fitting optimal ancestral palettes using badMIXTURE, on the residual scale shown. (E) Ancestral palettes estimated by badMIXTURE. 123 populations in total were simulated, with grey populations all being outgroups to those shown in colour.

In the “Ghost Admixture” scenario, P2 is instead formed by a 50%-50% admixture between P1 and an unsampled “ghost” population, which is most closely related to P3. For this scenario, the larger proportion of ancestry inferred from the light green population than the light pink one does not reflect a difference in admixture proportion, since neither P3 nor P4 actually contributed material to P2. Rather, it reflects the fact that P3 is more closely related to the unsampled ghost population, as seen in the phylogeny.

In the “Recent Bottleneck” scenario, P1 is a sister population to P2 that underwent a strong recent bottleneck. Members of P2 are once again inferred to be admixed, again with ancestry proportions that reflect phylogenetic distance rather than admixture proportions, while P1 receives its own ancestry component.

Since the “Ghost Admixture” and “Recent Bottleneck” scenarios cannot be represented using a simple admixture description, the model cannot be historically correct but the algorithm nevertheless attempts to fit the data as best it can by finding the combination of admixture proportions and ancestral frequencies that best explain the observed patterns. It is therefore useful in interpreting the results applied to real data to think of STRUCTURE and ADMIXTURE as algorithms that parsimoniously explain variation between individuals rather than as parametric models of divergence and admixture.

For example, if admixture events or genetic drift affect all members of the sample equally, then there is no variation between individuals for the model to explain. For example, non-African humans have a few percent Neanderthal ancestry, but this is invisible to STRUCTURE or ADMIXTURE since it does not result in differences in ancestry profile between individuals. The same reasoning helps to explain why for most datasets –even in species such as humans where mixing is commonplace - each of the *K* populations is inferred by STRUCTURE/ADMIXTURE to have non-admixed representatives in the sample. If every individual in a group is in fact admixed, then (with some technical caveats discussed by Falush et al. [4]) the model simply shifts the allele frequencies of the inferred ancestral population to reflect the fraction of admixture that is shared by all individuals.

The higher the value of *K* that is used, the less parsimonious the results of the model are as a description of the data. Specifically, each additional population allowed in a STRUCTURE or ADMIXTURE model requires many additional parameters to be inferred. First, every individual has a proportion of ancestry from the population that must be estimated. Secondly, every allele has an unknown frequency in the population. In common with other approaches, models with large numbers of parameters are algorithmically more difficult to fit to the data and also are penalized in statistical comparisons to prevent overfitting. Large numbers of parameters can lead to some undesirable algorithmic behaviour. For example, because the number of parameters increases with the number of loci, the algorithms can fail to detect subtle population structure in relatively simple scenarios even if the number of loci is very large [10].

A number of methods have been developed to estimate the number of populations *K* [1, 2, 11], which is a difficult statistical problem, even for data simulated according to the model assumptions, where there is by construction a true value to be inferred. For real data, the assumption that there is a true value is always incorrect; the question rather being whether the model is a good enough approximation to be practically useful. First, there may be close relatives in the sample which violates model assumptions [12]. Second, there might be “isolation by distance”, meaning that there are no discrete populations at all [13]. Third, population structure may be hierarchical, with subtle subdivisions nested within diverged groups. This kind of structure can be hard for the algorithms to detect and can lead to underestimation of *K* [14]. Fourth, population structure may be fluid between historical epochs, with multiple events and structures leaving signals in the data [15]. Many experienced users address these issues by examining the results of multiple *K* simultaneously. This makes interpreting the results more complex, especially because it makes it easier for users to find support for preconceptions about the data somewhere in the results.

In practice, the best that can be expected is that the algorithms choose the smallest number of ancestral populations that can explain the most salient variation in the data. Unless the demographic history of the sample is particularly simple, the value of *K* inferred according to any statistically sensible criterion is likely to be smaller than the number of distinct drift events that have significantly impacted the sample. In practice, the algorithm can use variation in admixture proportions between individuals to approximately mimic the effect of more than *K* distinct drift events without estimating ancestral populations corresponding to each one. In other words, an admixture model is almost always “wrong” (Assumption 2 of the Core protocol, Figure 1) and should not be interpreted without examining whether this lack of fit matters for a given question.

To be specific, in the Ghost Admixture scenario of Figure 2, the ghost population is modelled as being a mix of the sampled populations it is most closely related to, rather than being given its own ancestral population. In the Recent Bottleneck scenario, the genetic drift shared by P1 and P2 is modelled by ADMIXTURE by assigning both populations some ancestry from the light blue ancestral population. The strong recent drift specific to P1 is approximately modelled by assigning more light blue ancestry to P1 than to P2, thereby making P1 more distinct from the other populations in the sample. An alternative outcome in both scenarios would be for ADMIXTURE to infer a higher value of *K* and to include an extra ancestral population for P2. The algorithm is more likely to infer this solution if there was stronger genetic drift specific to P2 or if members of the population made up a greater overall proportion of the sample.

### Sample sizes matter

STRUCTURE/ADMIXTURE results are greatly affected by sample size [16]. Specifically, groups that are contain fewer samples or have undergone little population-specific drift of their own are likely to be fit as mixes of multiple drifted groups, rather than assigned to their own ancestral population. Indeed, if an ancient sample is put into a dataset of modern individuals, the ancient sample is typically represented as an admixture of the modern populations (e.g. [17], [18]), which can happen even if the individual sample is older than the split date of the modern populations and thus cannot be admixed.

The sensitivity of STRUCTURE/ADMIXTURE to sample size and to strong genetic drift requires the addition of “Algorithm input” (Step 0) to the protocol in Figure 1.

The problem of sampling strategy affecting inference is common to many methods. Principle Components Analysis (PCA) is closely related to the STRUCTURE model in the information that it uses, both in theory [10] and in practice [19] and has also been shown theoretically to be affected by sample size [6]. A neighbour-joining tree, calculated based on F_st_ or other measures of variation between populations also exaggerates the effect of drift, an example of which is shown in Figure 5G from Friedlaender et al, reproduced in Figure 3. Here, Africa and the Middle East together comprise a small part of the overall diversity, which is dominated by the isolated populations of Native Americans and Papua New Guinea as we discuss further in Case Study 2.

**Figure 3.**
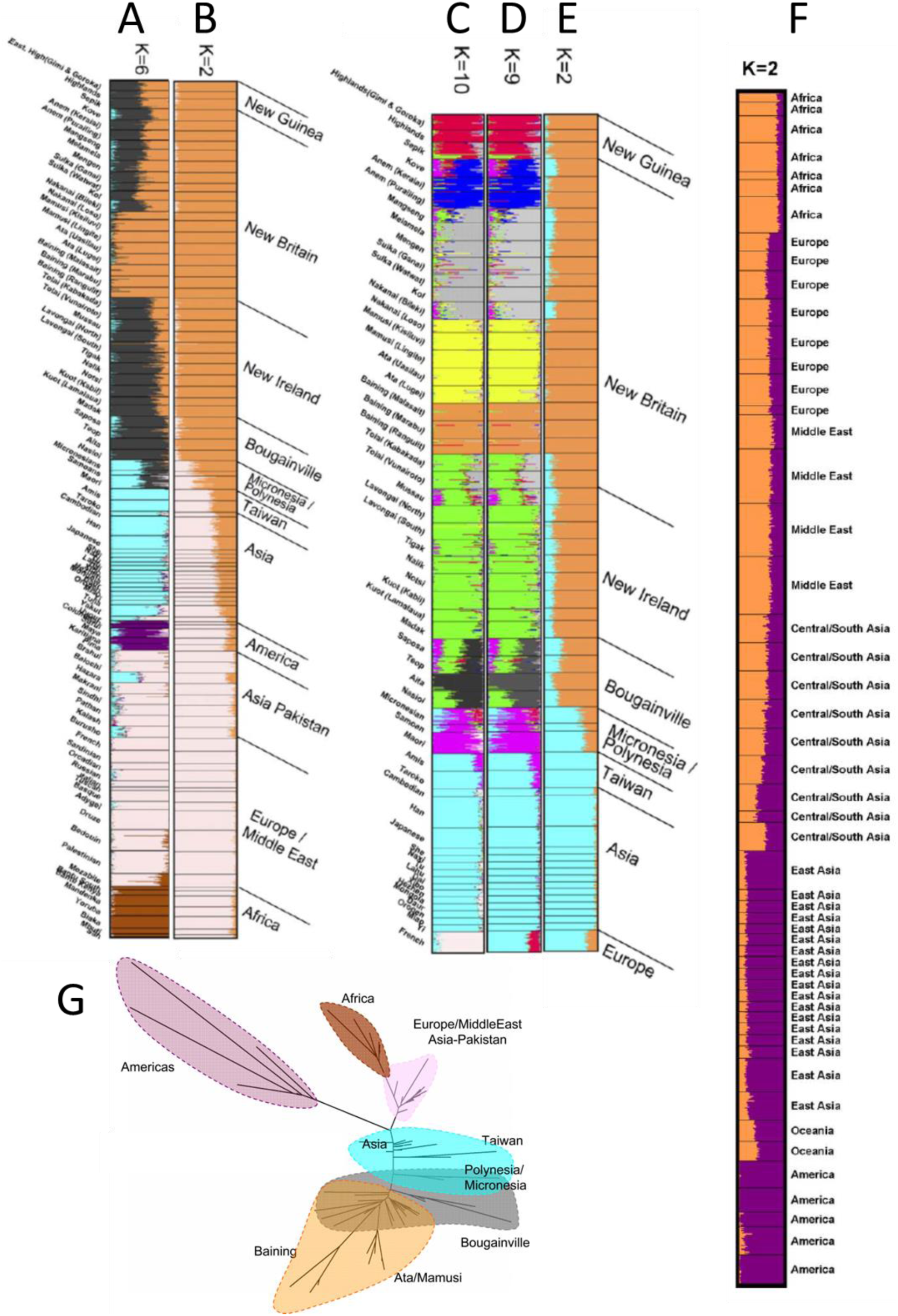
STRUCTURE results for global human genetic datasets reproduced from Friedlaender et al. [20] (A-E) and Rosenberg et al. [21] (F). (G) reproduces the neighbour-joining Fst tree [20] coloured according to K=6 STRUCTURE results (A).

**Figure 4.**
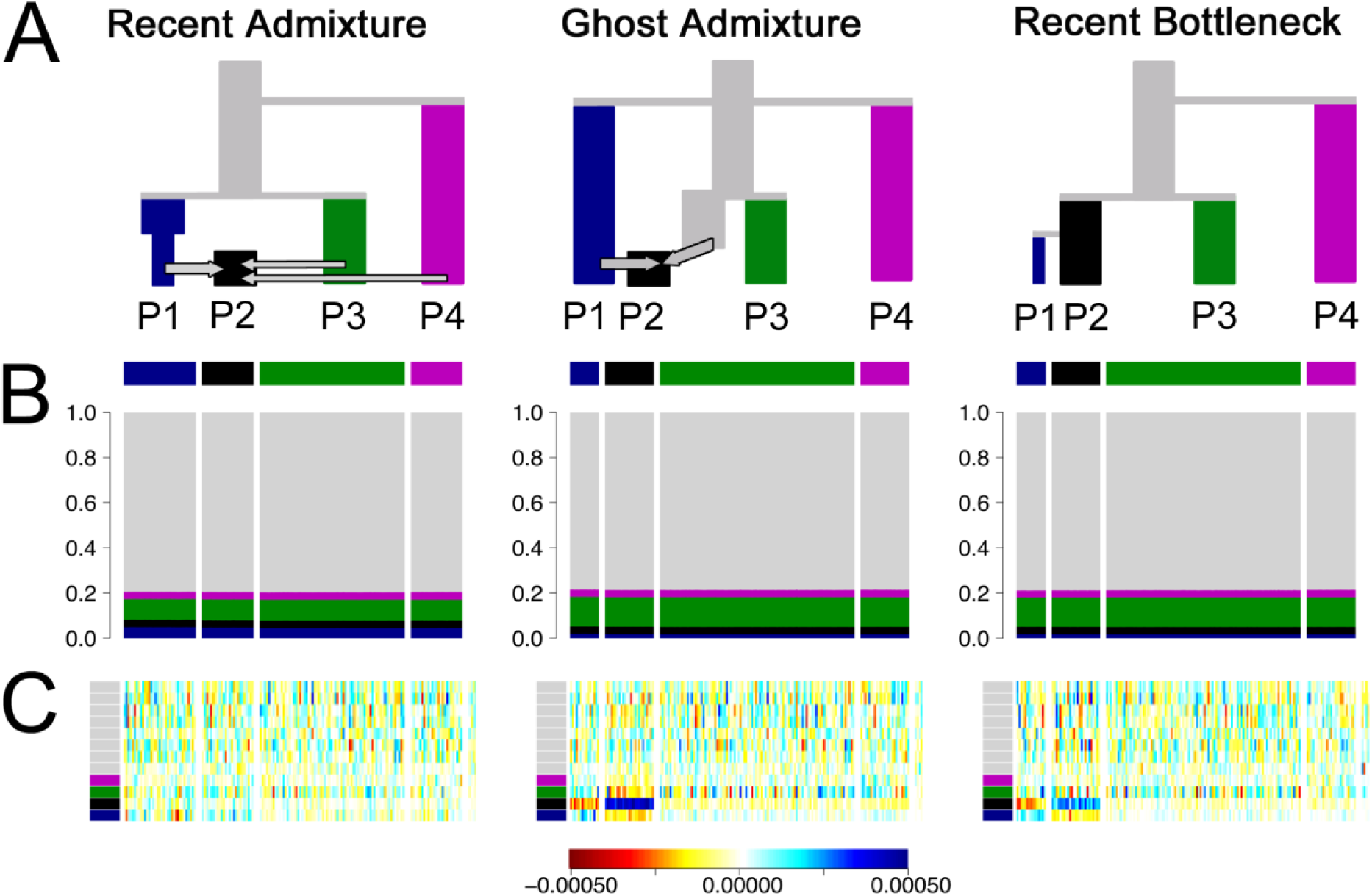
Unlinked badMIXTURE results for simulated data presented in Figure 2 using unlinked data under the same scenarios. Whilst the palettes look dramatically more homogeneous without linkage information (B vs Figure 2C), the badMIXTURE residuals (C vs Figure 2D) follow the same pattern, i.e. they are unstructured in the Recent Admixture data (scale shown below main plots).

**Figure 5.**
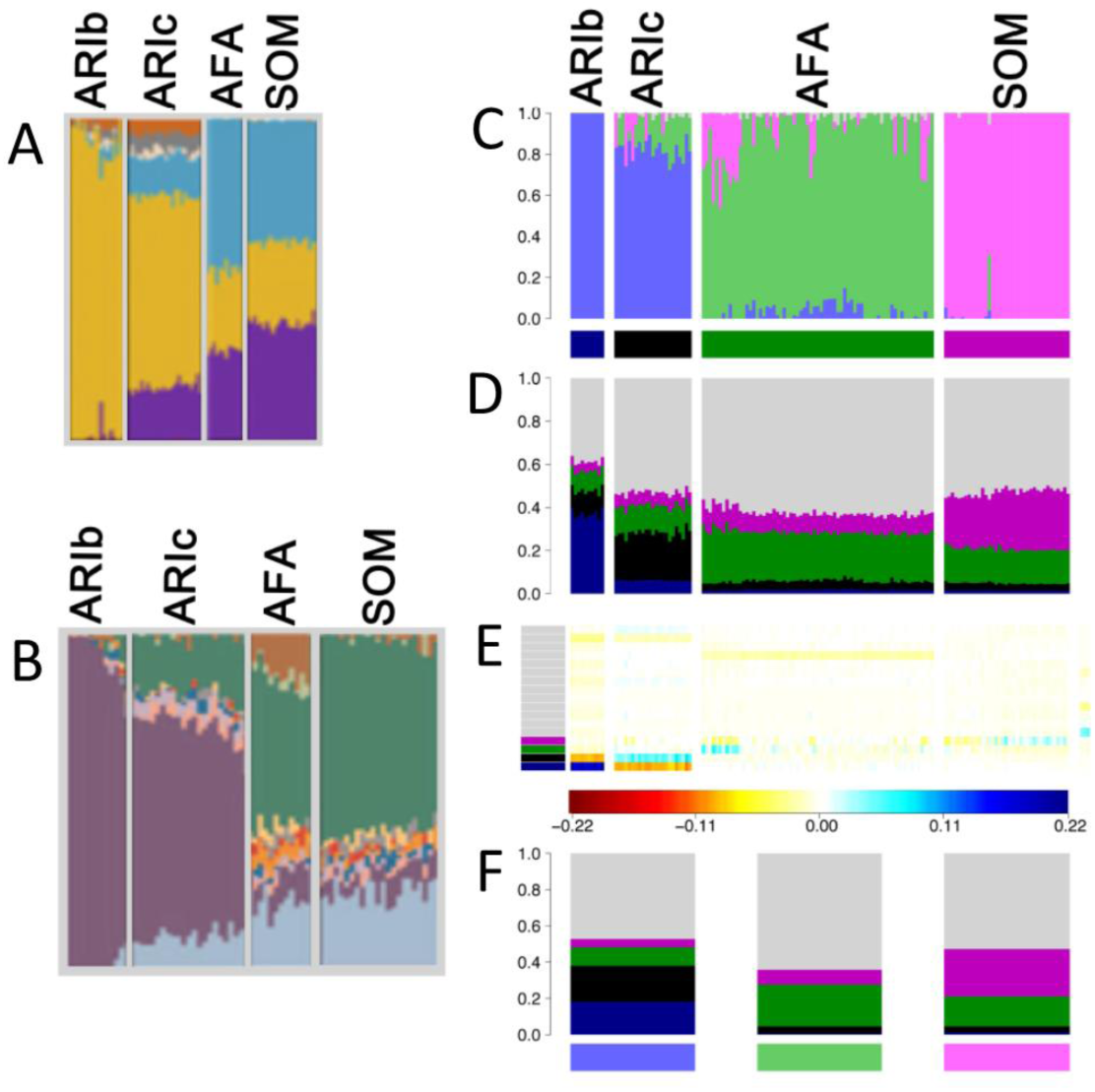
Analysis of Ari ancestry. ADMIXTURE analyses of the Ari and neighbouring Ethiopian groups adapted from (A) Pagani et al. [25], (B) Hodgson et al.[27] and (C) van Dorp et al. [26] at K = 11. Somali (SOM) and Afar (AFAR), Ari Blacksmith (ARIb) and Ari Cultivator (ARIc) populations were used in all three of the studies but the other populations differ substantially and the exact individuals differ slightly due to different quality control procedures and dataset merges. (D) CHROMOPAINTER inferred painting palettes based on (C). (E) badMIXTURE palette residuals under best fit ancestral population admixture model (F) Estimated ancestral palettes. Contributions from other populations are shown in grey.

#### Case Study 2: Worldwide human data

The effects of sample size are vividly illustrated by the analyses of Friedlaender et al. [20] who augmented a pre-existing microsatellite dataset from a worldwide collection by a similar number of samples from Melanesia, in order to study genetic relationships between Melanesians, for which purpose their sample was excellent. For *K*=2, their analysis infers Papua New Guinea (PNG) as one ancestral population and Western Eurasia and Africa as the other, with East Asians being represented as genetic mixtures (Figure 3B). This analysis differs from that of Rosenberg et al. [21] for *K*=2 who had only a small number of Melanesians in their sample, and who found Native Americans rather than Melanesians to be the unadmixed group (Figure 3F). For *K*=6, both models distinguish between all 5 continental groups (Americans, Western Eurasians, Africans, East Asians, and Oceanians), however Rosenberg et al. split Native American groups into two ancestral populations (not presented here), while Friedlaender et al. infer that Melanesians have two ancestral populations, with pure representatives in Bourganville and New Britain (Figure 3A). Rosenberg et al. [7] also found the Kalash, an isolated population in Pakistan, to be the sixth cluster.

It is tempting to attribute the global clustering results of Friedlaender et al. as being due to peculiar sampling but for *K*=2, the results of Rosenberg et al. [21] are actually odder, if interpreted literally, since they imply a continuous admixture cline between Africa and the Americas. From almost any perspective, the most important demographic event that has left a signature in the dataset is the out-of-Africa bottleneck. This is not taken by STRUCTURE to be the event at *K*=2 in either of the analyses, or that of others with similar datasets, because sub-Saharan Africans constitute only a small proportion of the sample.

Some even more peculiar results are obtained for an analysis that focused on Melanesian populations, leaving in only East Asian populations and a single European population, the French. Friedlander et al.’s purpose in presenting this analysis was to analyse the fine-scale relationships amongst the Melanesians, while detecting admixture e.g. from Colonial settlers. Our purpose here is to ask what the results imply, when interpreted literally, about the relationships between Melanesians, East Asians and Europeans. For all values from *K*=2 to *K*=9, the French population is inferred to be a mixture between an East Asian population and a Melanesian one. For *K*=7 to *K*=9, the model represents the European population as a mixture of an East Asian population and one from New Guinea (Figure 3D,E). Only for *K*=10 do the French form their own cluster and in this case they are inferred to have variable levels of admixture from East Asians (Figure 3C). Throughout, interpretation of the ancestral populations based on where individuals are today (Interpretation Protocol of Figure 1) would only make these results more misleading, implying at K=9 that the French are admixed between East Asians and Papuan highlanders.

Once again, it is tempting to write these results off as being the product of the sampling scheme, but the problem is fundamental to any approach based on equally weighing samples. If we instead imagine that there was an environmental catastrophe that spared the people of Melanesia and a few lucky others then the analysis would become a faithful sampling of the people of the world and the results would become the world’s genetic history and the literal interpretation of the bar plots would give misleading results, despite being proportionate sampling of extant humans.

This exercise is relevant in particular because human history is in fact full of episodes in which groups such as the Bantu in Africa, the Han in Asia, and the Northern Europeans in America have used technological, cultural or military advantage or virgin territory to multiply until they make up a substantial fraction of the world’s population. The history of the world told by STRUCTURE or ADMIXTURE is thus a tale that is skewed towards populations that are currently large and to those that have grown from small numbers of founders, with the bottlenecks that that implies. Even if the sampling is strictly proportional to modern population sizes, it is a winner’s history. In species that have not suffered environmental catastrophe, sampling choices are inevitable, are necessarily subjective and can greatly affect the clusters that are identified.

### badMIXTURE: a tool to visualise goodness of fit for admixture models

Experienced researchers, particularly those interested in population structure, typically present STRUCTURE results alongside other methods that make different modelling assumptions. These include TreeMix [22] ADMIXTUREGRAPH[23], fineSTRUCTURE [10], GLOBETROTTER [15], f3 and D statistics[24], amongst many others. These models can be used both to probe whether assumptions of the admixture model are likely to hold and to validate specific features of the results. Each also comes with its own pitfalls and difficulties of interpretation. It is not obvious that any single approach represents a direct replacement as a data summary tool, especially for datasets where recent mixture has caused inter-individual variation in admixture proportions.

Here, we build more directly on the results of STRUCTURE/ADMIXTURE by developing a new approach, badMIXTURE to examine which features of the data are poorly fit by the model, by looking at patterns within residuals. Rather than intending to replace more specific or sophisticated analyses, we hope to encourage their use by making the limitations of the initial analysis clearer.

An admixture model can only be a parsimonious way of describing the data if there are more distinct ancestry profiles than there are ancestral populations, since otherwise each ancestry profile could simply be assigned its own ancestral population. Therefore badMIXTURE assumes there are more distinct ancestry profiles *P* than there are populations *K*. In this article we use sampling labels to identify groups with distinct ancestry profiles, but if these are not available or are not predictive of genetic relationships, it is possible to use fineSTRUCTURE [10] to cluster individuals into genetically homogeneous groups based on their inferred palettes, thus generating labels.

badMIXTURE uses patterns of DNA sharing to assess the goodness of fit of a recent admixture model to the underlying genetic data. These sharing profiles are generated using CHROMOPAINTER [10] which calculates, for each individual, which of the other individual(s) in the sample are most closely related for each stretch of genome, using either haplotype or allele matching. This process is called “chromosome painting”, and can be thought of in terms of “palettes” (Figure 2C), which can also be visualized as barplots. The palette measures the proportion of the genome of each individual that is most closely related to the individuals sampled from each of the labelled populations. The painting palettes differ for the three simulated scenarios (Figure 2C), showing that there should be information in the genetic data to distinguish between them, even though they give almost identical ADMIXTURE barplots.

STRUCTURE and ADMIXTURE estimate both the ancestral gene frequencies and the admixture proportions for each individual in the sample. badMIXTURE assumes that the admixture proportions estimated by STRUCTURE and ADMIXURE are correct and uses matrix factorization to find the combination of ancestral palettes that give the best overall fit (evaluated using least squares) to the palettes of each individual. Crucially, under a number of reasonable assumptions (see Methods), in a recent admixture scenario, the palettes of admixed individuals should be a mixture of the palettes of non-admixed individuals according to the relevant admixture proportions.

In other words, if a simple admixture scenario is correct and the proportions are correctly estimated by STRUCTURE/ADMIXTURE, then it should be possible to use the *N* × *K* admixture proportions of the *N* individuals in the sample and the *K* × *P* palettes proportions for the *K* ancestral populations to predict the *N* × *P* palette proportions for each individual. The fit of the model can be examined by comparing the true palette proportions for each individuals to the ones predicted by badMIXTURE.

Figure 2D shows the residuals, representing the difference between the observed palettes for each individual in the simulated data and those reconstructed by badMIXTURE. Figure 2E shows the corresponding palettes inferred for each ancestral population. Under the Recent Admixture scenario, there is no systematic pattern to the residuals. For the Ghost Admixture scenario, the residuals show a systematic pattern, with the model substantially underestimating the proportion of palette that individuals in P2 have from their own population and overestimating the contributions from the other populations. For the recent bottleneck model, the deviations are similar - the main qualitative difference between the Ghost Admixture scenario and Recent Bottleneck scenario are in the ancestral palettes. Ghost admixture produces a much more uniform ancestral palette than either of the other models, which both contain bottlenecks for P1.

badMIXTURE distinguishes the Recent Admixture scenario from alternatives because the recent admixture model makes the distinctive prediction that admixed individuals are not particularly related to each other, as shown by the small amount of black in their palettes in Figure 2C. Members of P2 get 50% of their genomes from the light blue ancestral population, 35% from the light green population and 15% from the light pink one, while P1 received all of its ancestry from the light blue population. For any given locus, a member of P2 will have the same ancestral source as a member of P1 50% of the time. But two members of P2 will have the same ancestry source only 0.5^2^ + 0.35^2^ + 0.15^2^ = 0.395 of the time. This means that paradoxically, members of P2 may (depending on the exact details of population history) be more related to members of P1 than they are to each other and have relatively little of their palette from their own population. Under the other scenarios, individuals from P2 receive more of their palette from other members of their own population.

### Goodness of fit aids interpretation even without linkage information

STRUCTURE/ADMIXTURE has been applied to thousands of different species, most of which do not have the linkage maps (either physical or genetic) required for chromosome painting. The algorithm can also be applied to datasets with relatively small number of markers. It would therefore be advantageous to be able to apply a similar approach to these datasets.

An equivalent analysis can be performed using chromosome painting in unlinked model, as shown in Figure 4, to generate allele-sharing palettes. The results are qualitatively similar to the CHROMOPAINTER analysis exploiting Linkage Disequilibrium; however because the palettes are closer to uniform (Figure 4B), the residuals contain more noise (Figure 4C). If few markers were available, there may be no interpretable signal remaining making it impossible to distinguish between different scenarios on the basis of limited genetic information.

#### Case study 3: The Ari of Ethiopia

This example highlights a situation in which application of badMIXTURE could have prevented a false history from being inferred. Three sets of researchers [25–27] investigated the relationships between the origins of occupational groups (Blacksmiths and Cultivators) in the Ari community of Ethiopia, all applying ADMIXTURE analyses (Figures 5A, B, C). The first two sets of researchers tentatively concluded that the two groups were most likely to have had different ancestral sources.

First, Pagani et al. [25] analysed the data and cautiously interpreted the ADMIXTURE results:

*One insight provided by the ADMIXTURE plot (Figure [5A]) concerns the origin of the Ari Blacksmiths. This population is one of the occupational caste-like groups present in many Ethiopian societies that have traditionally been explained as either remnants of hunter-gatherer groups assimilated by the expansion of farmers in the Neolithic period or as groups marginalized in agriculturalist communities due to their craft skills. The prevalence of an Ethiopian-specific cluster (yellow in Figure [5A]) in the Ari Blacksmith sample could favor the former scenario; the ancestors of this occupational group could have been part of a population that inhabited the area before the spread of agriculturalists.*

This interpretation was supported by a similar analysis by Hodgson et al. [27]:

*As the Ari Blacksmiths have negligible EthioSomali ancestry, it seems most likely that the Ari Cultivators are the descendants of a more recent admixture between a population like the Ari Blacksmiths and some other [Horn Of African] population (i.e. the Ethio-Somali ancestry in the Ari Cultivators is likely to substantially postdate the initial entry of this ancestry into the region).*

van Dorp et al [26] found similar ADMIXTURE results. Interpreted according to the protocol above, these analyses all imply that the Blacksmiths are pure representatives of one ancestral population (as shown by a homogeneous block of colour), while Cultivators are recently admixed, receiving ancestral contributions from neighbouring Ethiopian groups. However, the results of the three studies have different sampling and differ in how much of the ancestral population that Blacksmiths purportedly represent has contributed to the Cultivators or to other groups.

However, van Dorp et al [26] used additional analyses including Ghost and Recent Bottleneck simulations, as in Figure 2, together with fineSTRUCTURE [10], and GLOBETROTTER [15] to show that this history is false and the totality of evidence from the genetic data supports that the true history is analogous to the Recent Bottleneck scenario. The Blacksmiths and the Cultivators diverged from each other, principally by a bottleneck in the Blacksmiths, which was likely a consequence of their marginalised status. Once this drift is accounted for the Blacksmiths and Cultivators have almost identical inferred ancestry profiles and admixture histories. In our analysis, a strong deviation from a simple admixture model can be seen in the residual palettes, which imply that the ancestral palettes estimated by badMIXTURE substantially underestimate the drift in the Ari Blacksmiths (Figure 5E).

#### Case study 4: Ancient Indian populations

Population genetic inference is hard. History is complex and consists of multiple discrete or continuous events happening in the near and distant past. Researchers may be interested in the distant past, while the strongest signals in the data often reflect more recent migrations. Models that attempt to account for complexity can often be unwieldy and it is difficult to infer large numbers of parameters at once. STRUCTURE and ADMIXTURE are popular because they give the user a broad-brush view of variation in the data, while also potentially allowing the possibility of zooming down on details about specific individuals or labelled groups. Ideally, statistical genetic methods would be developed that emulate these features, without suffering from the various difficulties of interpretation described above.

In our final example, we attempt to address the challenge of complex inference by providing an overview of a history in a single figure. As in previous case studies, the approach we take here is to build directly on the clustering results of ADMIXTURE from a published example and evaluate their fit using badMIXTURE. Once again, it is important to emphasize the limitations inherent to this approach, which should be addressed by testing specific inferences of interest using other methods.

Basu et al. [28] used an ADMIXTURE plot with K=4 to summarize variation amongst continental Indians from 19 labelled groups. The four ancestral populations were labelled Ancestral North Indian (ANI), Ancestral South Indian (ASI), Ancestral Tibeto-Burman (ATB) and Ancestral Austro-Asiatic (AAA), as shown in Figure 6A. They argued that a major conclusion from their analysis is that the structure of mainland India is best described by 4 ancestral components.

**Figure 6.**
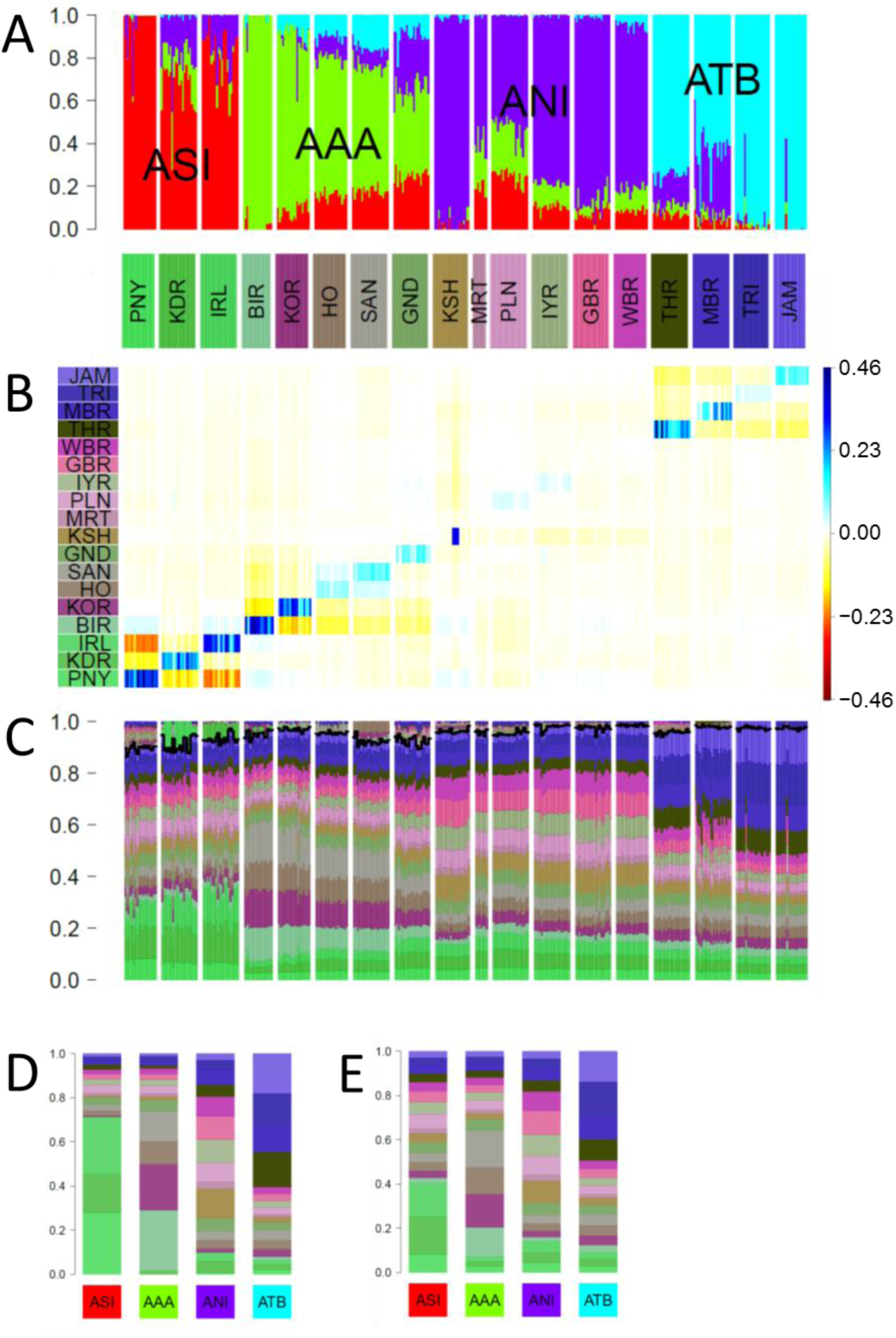
Comparison of ADMIXTURE with painting palettes for Indian genetic data originally presented in [28]. (A) ADMIXTURE profile at K = 4 (B) Residuals palettes estimated by badMIXTURE. (C) Painting palettes after correcting within-population values as described in text. The part of the palette above the black line is not predicted by badMIXTURE (D) Ancestral palettes estimated by badMIXTURE (E) Estimated ancestral palettes after correcting for within-population values.

The overall fit of the ADMIXTURE results estimated by badMIXTURE is poor (Figure 6B). However, the large residuals are primarily *within* ancestral components, i.e. structured in a block-diagonal form, with blocks that correspond to the four ancestral components estimated by ADMIXTURE. Furthermore, almost all of the positive residuals are on the diagonal, i.e. specific to the labelled group that the individual was assigned to. These residuals vary substantially according to ancestral component, with ASI populations having the highest on-diagonal residuals and the ANI populations having the lowest ones. Within the KSH (Kharti) population, there is substantial variation amongst individuals, presumably reflecting the presence of relatives or other strong sub-structure within the labelled group.

The structure of these residuals suggests that they principally reflect recent genetic drift that is specific to labelled groups, with considerable variation amongst groups in how much drift has occurred, presumably reflecting their recent demographic history. However, the block-like form suggests that if this recent genetic drift can be accounted for, the data might still be consistent with a history of mixture of four ancestral components, as suggested by the initial ADMIXTURE results.

We have implemented a simple procedure within badMIXTURE to estimate the composition of painting palettes in the absence of group specific drift (see Methods), which for this dataset substantially reduces the residuals (not shown). Figure 6C shows the corrected individual palettes, and the ancestral palettes are substantially altered by the removal of recent drift, particularly for ASI and AAA populations (compare Figure 6D-E). A more rigorous but laborious approach to removing label-specific drift, namely to remove individuals with the same label from the donor panel used for chromosome painting, was implemented by van Dorp et al. [26].

Examining the corrected palettes shown in Figure 6C carefully, it is possible to see evidence that there are indeed four distinct ancestral components in the data, validating the major claim made by Basu et al. in their original analysis. For example, the three labelled groups with high ASI ancestry have similar palettes that are clearly distinct from those of all other labelled groups, with all of them having large amounts of green of three different shades. However, comparison of these palettes with the ADMIXTURE results also highlights the likely effect of recent genetic drift on those results, which is analogous to but less dramatic than that observed in the Ari case study. Specifically, the PNY (Paniya) are inferred by ADMIXTURE to be the only unadmixed representatives of the ASI population (Fig 6A) but also have the largest badMIXTURE residuals prior to correction, which presumably reflects recent drift (Fig 6B). After correction, PNY actually receive a smaller proportion of their palette from ASI groups than the other two ASI groups do (Fig 6C).

This analysis highlights the fact that the mixture fractions estimated by ADMIXTURE are unreliable and that no individual group can be safely assumed to be pure representatives of the ancestral source. That said, the relative admixture fractions are more plausible for the other three ancestry components, since the labelled groups that are estimated as pure by ADMIXTURE, i.e. BIR (Birhor), KSH and TRI (Tripuri)/JAM (Jamatia) also have the highest contribution from ancestrally related groups within their painting palettes.

Careful examination of these palettes also provide evidence of sharing of ancestry between pairs of populations that is not predicted based on the four ancestral palettes (shown above the black line in Figure 6C), providing further evidence of the importance of recent demography, rather than ancestral population mixture in shaping diversity. These pairs of populations are TRI and JAM, IRL (Irula) and KDR (Kadar), HO (Ho) and SAN (Santal) and BIR and KOR (Korwa). This sharing is most likely to have arisen during the divergence of the populations from each other. This might be due to shared drift or recent patterns of migration.

Overall, this analysis provides evidence for demographic events at multiple scales. At the most local and recent scales, there is evidence for heterogeneity within groups, as shown by individuals within labelled groups with atypical ADMIXTURE profiles (e.g. in TRI and JAM) or badMIXTURE residuals (e.g. KSH). This heterogeneity provides evidence for recent migration between groups and substructure within groups, respectively. However, with the exception of some of the ANI populations, individuals in each group are distinguishable according to their badMIXTURE residuals and also based on fineSTRUCTURE clustering (see supplementary information of Basu et al.). This shows that most of the labelled groups are samples from populations that have been distinct from each other for long enough to acquire distinct and distinguishable genetic identities.

At the largest and most ancient scale, there is evidence for four ancestry components with clearly distinct painting palettes. However, the analysis in itself provides little evidence about the origin of these four ancestry components and the processes that gave rise to them, which would be best elucidated by relating the diversity found in India to that found in a global reference panel and by demographic modelling.

The greatest challenges in model interpretation occur at intermediate scales. There is clear evidence for admixture between ancestry components in some of the populations, such as GND (Gond) and MBR (Manipuri Brahmin). However, interpretation is made harder by the effects of recent genetic drift on ADMIXTURE estimates, seen most clearly for ASI populations. There is also evidence of shared ancestry between pairs of groups that is not predicted by the four component model, showing that recent patterns of migration between regions have played a significant part in shaping genetic diversity. However, a much more involved analysis would be necessary to elucidate these migration patterns and to relate them to the overall patterns observed.

### Conclusions

Notwithstanding the pitfalls we have described, there is tremendous value in summarizing data based on a handful of major axes of variation. Identifying these axes is a first step in historical reconstruction and in asking whether they can in fact be related to specific historical bottlenecks, expansions or migrations or whether they instead reflect continuously acting processes. PCA is effective for representing two or three axes of variation but becomes unwieldy for four or more and suffers from similar interpretation issues as STRUCTURE/ADMIXTURE, without the benefit of having an explicit model and hence model fit to check.

It is rarely the case that sampled data follows a simple history comprising a differentiation phase followed by a mixture phase, as assumed in an admixture model and highlighted by case study 1. Naïve inferences based on this model (the Protocol of Figure 1) can be misleading if sampling or choice of *K* are inappropriate, or if recent bottlenecks or unobserved ancient structure appear in the data. We have provided a tool, badMIXTURE, that identifies when this is likely to be the case.

The popularity of STRUCTURE and its descendants as unsupervised clustering methods means that they will be applied and interpreted, for which badMIXTURE provides important assistance. However, these analyses should always be followed up with tests of specific hypotheses, using other approaches. Running STRUCTURE or ADMIXTURE is the beginning of a detailed demographic and historical analysis, not the end.

## Materials and Methods

### Simulations

Figure 2A illustrates the demographic histories behind three simulation scenarios we name “Recent Admixture”, “Ghost Admixture” and “Recent Bottleneck”.

These simulations comprise a subset of full simulations described in [26] which aim to capture global human population genetic diversity across 13 simulated world-wide populations. Here, for tractability and motivated by Case Study 3, we explore the impact of different demographic histories in a subset of simulated groups: P1-P4. The simulation protocol used to generate the world-wide out-group populations is described in full detail in Supplementary Note 1.

For the “Recent Bottleneck” and “Ghost Admixture” simulations 13 populations were simulated using the approximate coalescence simulation software MaCS [29] under histories that differ in how P2 relates to P1 (Figure 2A). In the “Recent Bottleneck” P1 splits from P2 20 generations ago followed immediately by a strong bottleneck in P2. In the “Ghost Admixture” scenario, instead P1 splits from P2 1700 generations ago after which migrants from P1 form approximately 50% of P2 over a period of 200-300 generations. Although simulating 100 individuals in each population, we perform subsequent ADMIXTURE and CHROMOPAINTER analyses on a subset of these using only 35 individuals from P1, 25 individuals from P2, 70 individuals from P3 and 25 individuals from P4. This leaves an ‘excess’ of simulated individuals. For ease of interpretation only P1-P4 are depicted in Figure 2 with all out-group populations coloured grey.

For the “Recent Admixture” scenario we implement a simulation technique adapted from that applied in [30], related to that in [31], which sub-samples chromosomes from the ‘excess’ individuals simulated under the “Recent Bottleneck” scenario. This method explicitly mixes chromosomes from different populations based on a set of user-defined proportions, analogous to an instantaneous admixture event. Importantly for our purposes, this allows direct assessment of how well ADMIXTURE recapitulates these proportions. Using this approach we simulate admixed chromosomes of P2 by mixing chromosomes of 20 ‘excess’ individuals from each of P1 (50%), P3 (35%) and P4 (15%) based on an admixture event occurring λ=15 generations ago. In particular, to simulate a haploid admixed chromosome and as in Leslie et al. [30] we first sample a genetic distance *x* from an exponential distribution with rate 0.15 (λ/100). The first *x* cM of the simulated chromosome is composed of the first *x* cM of chromosomes selected randomly, but without overlap, from ‘excess’ individuals of P1, P3 and P4 according to the defined proportions. This process is repeated using a new genetic distance sampled from the same exponential distribution (rate=0.15) and continued until an entire simulated chromosome is generated. The method is then re-employed to generate a set of 20 haploid chromosomes for a single individual and then repeated 70 times to generate 70 haploid autosomes. Diploid individuals are constructed by joining two full sets of haploid chromosomes, resulting in 35 simulated P2 individuals in total.

#### Estimation of ADMIXTURE barplots and CHROMOPAINTER palettes

For each simulation scenario we apply ADMIXTURE [2] to the sampled individuals from every simulated group. SNPs were first pruned to remove those in high linkage disequilibrium (LD) using PLINK v1.07 [32] so that no two SNPs within 250kb have a squared correlation coefficient (r^2^) greater than 0.1. ADMIXTURE was then run with default values for multiple values of K, and the resultant admixture profiles plotted where K=11 (Figure 2B and Figure 4C). In addition for each scenario we applied CHROMOPAINTER to paint all individuals in relation to all others using default values for the CHROMOPAINTER mutation/emission (“-M” switch) and switch (“-n” switch) rates. When running CHROMOPAINTER ignoring information from Linkage Disequilibrium we use the unlinked mode (“-u” switch). We sum the total proportion of genome-wide DNA (linked) or matching chunk counts (unlinked) each recipient individual is painted by each donor group and plot the inferred contributions for each recipient as a painting palette.

### Estimation of ancestral palettes

Define *A* as the *N* × *K* admixture proportion matrix, where there are N individuals in the sample and K ancestral populations used in the ADMIXTURE analysis. Let *C* be the *N* × *P* matrix of individual palettes from the CHROMOPAINTER painting, and *X* be the *K* × *P* matrix of the palettes for each ancestral population. Then we seek solutions for *X* that minimise the squared prediction error of the form:

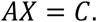

We define *B* = (*A*^*T*^*A*)^−1^*A*^*T*^. Then, *BAX* = (*A*^*T*^*A*)^−1^*A*^*T*^*AX* = *X*, leading to the solution

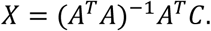

Note that there is no guarantee that *X* will be positive. Negative elements would imply a poor fit of the admixture model, and alternative minimization strategies might be employed to find *X* subject to the constraint. Further, if the matrix *A*^*T*^*A* is rank deficient its inverse will not exist. This should only be the case if *K* is chosen too large, or there are genuine symmetries in the data.

For a recent admixture model, long haplotypes are inherited from each of the donating populations in a given admixture proportion. If we assume that ancestral boundaries can be inferred then, excluding drift in either SNP frequency or haplotype structure, the palettes of admixed individuals are (by definition) a mixture with the same ancestry proportions as the SNPS under which admixture is inferred.

## Acknowledgements

This paper was stimulated by a discussion with Chuck Langley and also by the 2016 Workshop for Population and Speciation Genomics, Cesky Krumlov. We thank Jonathan Pritchard for suggesting the residuals plot and Analabha Basu, Kimberly Gilbert, Matthew Hahn, Razib Khan, Partha Majumder, Iain Mathieson, and Matthew Stephens for comments. D.F. is funded by a Medical Research Council Fellowship as part of the MRC CLIMB consortium for microbial bioinformatics (grant number MR/M501608/1). LvD is supported by the Newton Trust (MR/ P007597/1). DJL is supported by Wellcome Trust and Royal Society grant WT104125AIA.

## References

1. Pritchard, J.K., M. Stephens, and P. Donnelly, Inference of population structure using multilocus genotype data. Genetics, 2000. 155(2): p. 945–959.

2. Alexander, D.H., J. Novembre, and K. Lange, Fast model-based estimation of ancestry in unrelated individuals. Genome Research, 2009. 19(9): p. 1655–1664.

3. Tang, H., et al., Estimation of individual admixture: Analytical and study design considerations. Genetic Epidemiology, 2005. 28(4): p. 289–301.

4. Falush, D., M. Stephens, and J.K. Pritchard, Inference of population structure using multilocus genotype data: Linked loci and correlated allele frequencies. Genetics, 2003. 164(4): p. 1567–1587.

5. Novembre, J., Pritchard, Stephens, and Donnelly on Population Structure. Genetics, 2016. 204(2): p. 391–393.

6. McVean, G., A Genealogical Interpretation of Principal Components Analysis. Plos Genetics, 2009. 5(10).

7. Rosenberg, N., et al., Genetic structure of human populations. Science, 2002. 298(5602): p. 2381–2385.

8. Tishkoff, S., et al., The Genetic Structure and History of Africans and African Americans. Science, 2009. 324(5930): p. 1035–1044.

9. Rosenberg, N., et al., Empirical evaluation of genetic clustering methods using multilocus genotypes from 20 chicken breeds. Genetics, 2001. 159(2): p. 699–713.

10. Lawson, D.J., et al., Inference of Population Structure using Dense Haplotype Data. Plos Genetics, 2012. 8(1).

11. Evanno, G., S. Regnaut, and J. Goudet, Detecting the number of clusters of individuals using the software STRUCTURE: a simulation study. Molecular Ecology, 2005. 14(8): p. 2611–2620.

12. Anderson, E.C. and K.K. Dunham, The influence of family groups on inferences made with the program Structure. Molecular Ecology Resources, 2008. 8(6): p. 1219–1229.

13. Frantz, A.C., et al., Using spatial Bayesian methods to determine the genetic structure of a continuously distributed population: clusters or isolation by distance? Journal of Applied Ecology, 2009. 46(2): p. 493–505.

14. Janes, J.K., et al., The K=2 conundrum. Molecular Ecology, 2017. 26(14): p. 3594–3602.

15. Hellenthal, G., et al., A Genetic Atlas of Human Admixture History. Science, 2014. 343(6172): p. 747–751.

16. Puechmaille, S., The program structure does not reliably recover the correct population structure when sampling is uneven: subsampling and new estimators alleviate the problem. Molecular Ecology Resources, 2016. 16(3): p. 608–627.

17. Rasmussen, M., et al., Ancient human genome sequence of an extinct Palaeo-Eskimo. Nature, 2010. 463(7282): p. 757–762.

18. Skoglund, P., et al., Origins and Genetic Legacy of Neolithic Farmers and Hunter-Gatherers in Europe. Science, 2012. 336(6080): p. 466–469.

19. Patterson, N., A. Price, and D. Reich, Population structure and eigenanalysis. Plos Genetics, 2006. 2(12): p. 2074–2093.

20. Friedlaender, J., et al., The genetic structure of Pacific islanders. Plos Genetics, 2008. 4(1).

21. Rosenberg, N., et al., Clines, clusters, and the effect of study design on the inference of human population structure. Plos Genetics, 2005. 1(6): p. 660–671.

22. Pickrell, J.K. and J.K. Pritchard, Inference of Population Splits and Mixtures from Genome-Wide Allele Frequency Data. Plos Genetics, 2012. 8(11): p. 17.

23. Leppala, K., S.V. Nielsen, and T. Mailund, admixturegraph: an R package for admixture graph manipulation and fitting. Bioinformatics, 2017. 33(11): p. 1738–1740.

24. Patterson, N., et al., Ancient Admixture in Human History. Genetics, 2012. 192(3): p. 1065-+.

25. Pagani, L., et al., Ethiopian Genetic Diversity Reveals Linguistic Stratification and Complex Influences on the Ethiopian Gene Pool. American Journal of Human Genetics, 2012. 91(1): p. 83–96.

26. van Dorp, L., et al., Evidence for a Common Origin of Blacksmiths and Cultivators in the Ethiopian Ari within the Last 4500 Years: Lessons for Clustering-Based Inference. Plos Genetics, 2015. 11(8): p. 49.

27. Hodgson, J.A., et al., Early Back-to-Africa Migration into the Horn of Africa. Plos Genetics, 2014. 10(6): p. 18.

28. Basu, A., N. Sarkar-Roy, and P. Majumder, Genomic reconstruction of the history of extant populations of India reveals five distinct ancestral components and a complex structure. Proceedings of the National Academy of Sciences of the United States of America, 2016. 113(6): p. 1594–1599.

29. Chen, G., P. Marjoram, and J. Wall, Fast and flexible simulation of DNA sequence data. Genome Research, 2009. 19(1): p. 136–142.

30. Leslie, S., et al., The fine-scale genetic structure of the British population. Nature, 2015. 519(7543): p. 309-+.

31. Price, A.L., et al., Sensitive Detection of Chromosomal Segments of Distinct Ancestry in Admixed Populations. Plos Genetics, 2009. 5(6).

32. Purcell, S., et al., PLINK: A tool set for whole-genome association and population-based linkage analyses. American Journal of Human Genetics, 2007. 81(3): p. 559–575.

