## Supplementary Materials for "A tutorial on how (not) to over-interpret STRUCTURE/ADMIXTURE bar plots"

**Supplementary Note 1: Full Simulation Protocol**

Overview

Simulations were designed to approximately mimic human population history and have strong similarities to the simulation scenarios described in Hellenthal et al^1^ and van Dorp et al^2^. As also noted in these publications, it is impossible to simulate a scenario that perfectly captures the history of modern human groups, reinforced by references in the literature on appropriate split times and population sizes often uncertain or disagreeing, however here we aim to capture major features of world-wide human migrations and splits, informed by a number of publications ^3–7^.

In the context of this work and for simplicity we mainly consider the impact of the simulated demographic history on inferences for 4 populations: P1-P4, representing Ethiopian-like populations in Figure S1. This is motivated by both the focus on Ethiopian populations in main text Case Study 3, as well as the use of predominately human examples throughout the manuscript which require evaluation of complex global histories. For completeness here we describe the simulation procedures in full.

Simulation Method

Simulations were performed using the approximate coalescence simulation software MaCS^8^ with 13 populations simulated in total. In all cases simulations were conducted to produce 20 independent regions of size 64Mb each, but with variation in recombination rates based on HapMap Phase 2 build 36 genetic maps for chromosomes 1-20, respectively. Assuming a genome-wide average recombination rate of 1.25 × 10−8 and a mutation rate of 4 × 10−8 per basepair per generation, 200 haplotypes (100 individuals) were sampled from each of the 13 populations. For each of the 20 regions, 14,225 SNPs were selected (284,500 SNPs in total) such that the minor allele frequency spectrum of the simulated dataset (across all populations) matched that of the global dataset used in van Dorp et al 2015 across 100 equally spaced bins from 0 to 0.5. In downstream ADMIXTURE and CHROMOPAINTER analyses, of the 100 simulated individuals per population, we vary the numbers of individuals sampled from each population as follows: Afr1 (*n*=25), Afr2 (*n*=75), Afr3 (*n*=95), Afr4 (*n*=60), Eur1(*n*=100), Eur2 (*n*=100), MidE1 (*n*=0), EA1 (*n*=100), EA2 (*n*=75), P1 (*n*=35), P2 (*n*=25), P3 (*n*=70), P4 (*n*=25).

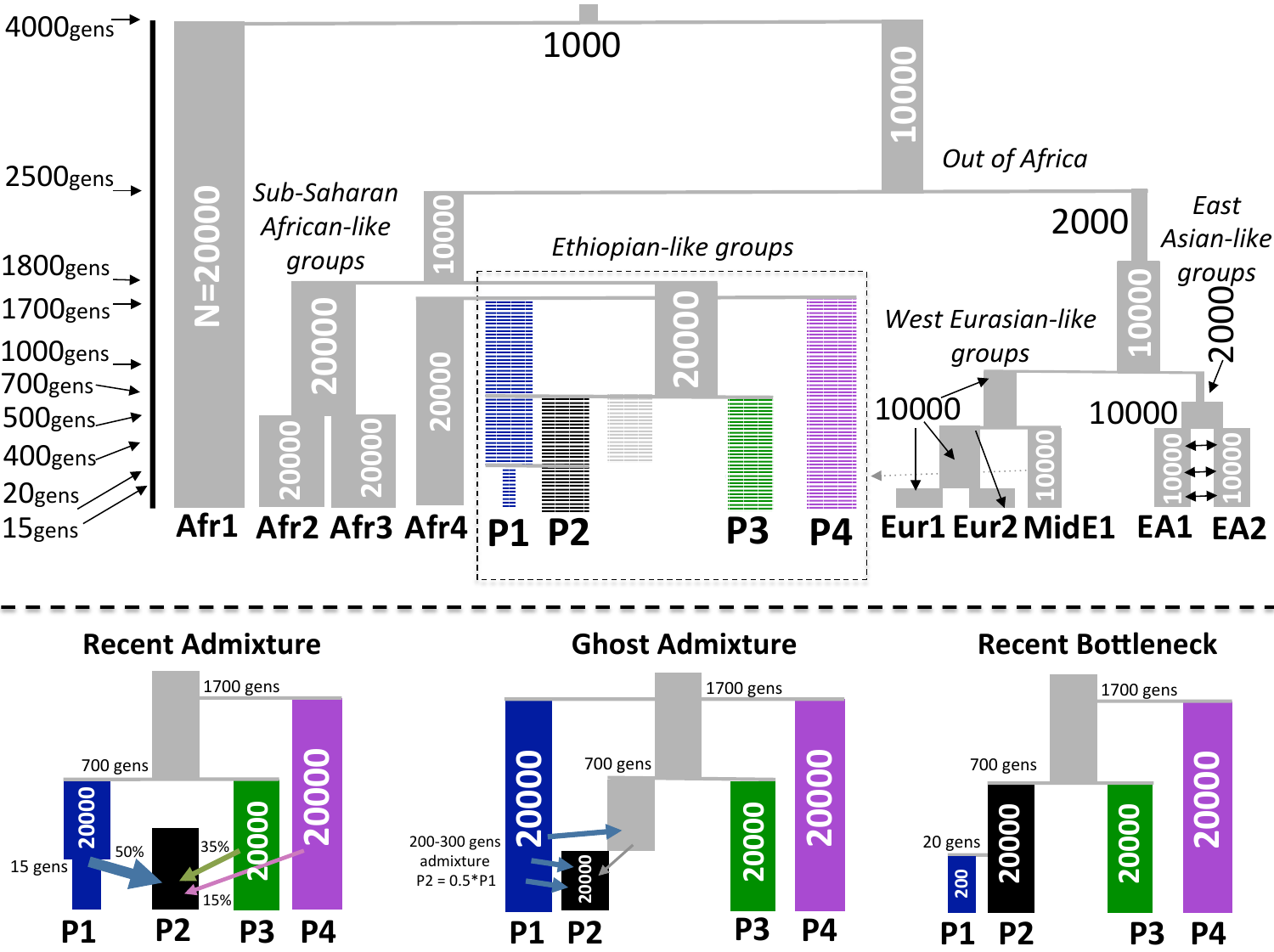

Figure S1 (top) Full simulation scenario of 13 populations designed to approximately mimic human demographic history. The time-line at left gives the approximate split times as also described in the text. The numbers within population bars provide the effective population sizes used in the simulations. The black dashed box highlights the Ethiopian-like populations P1-P4, which are the groups considered explicitly in this work and explored in eg. main text Figure 2 and Figure 4. (bottom) Demographic histories of populations P1-P4 under the Recent Admixture, Ghost Admixture and Recent Bottleneck simulation scenarios.

Global simulation scenario

Under all simulations Afr1-Afr4 and P1-P4 of Figure S1 are designed to mimic genetic diversity among African populations^6^. The split at 2,500 generations ago and subsequent bottleneck in simulated non-African populations Eur1-Eur2, MidE1 (European and Middle Eastern like groups) and EA1-EA2 (East Asian like groups) is designed to mimic the Out-of-Africa event^3,6,7^, and the split between Eur1-Eur2, MidE1 and EA1-EA2 at 1,000 generations ago mimics the split between Western Eurasian and East Asian populations ^4,5^ with a bottleneck in the latter. Continuous symmetric migration between EA1 and EA2 was also included such that each population’s fraction of new migrants increased by 0.00025 each generation for 375 gens (continuing until present- day), in order to represent a scenario of stable long-term migration between nearby populations.

For all simulation scenarios, 100 generations denotes the split between European-like groups Eur1 and Eur2; 375 generations the split between East Asian like groups EA1 and EA2; 400 gens the split of Eur1-Eur2 from Middle-East like group MidE1; 500 gens the split of African groups Afr2 and Afr3; 700 gens the split of Eur1/Eur2/MidE1 and EA1/EA2; 1,800 gens the split of Afr2 and Afr3 from Afr4 and P1-P4; 2,500 gens the split of Afr2-Afr4, P1-P4 from Eur1-Eur2, MidE1, EA1-EA2 and 4,000 gens the split of Afr1 from all other populations.

Additionally to represent proposed admixture from outside of Africa (back to Africa)^9^, the simulation included one-way admixture from an un-sampled Middle Eastern population (MidE1) into P1-P4, such that the fraction of new migrants into these groups increases by 0.02 for 10 gens, beginning 100 generations ago. Thus the total proportion of admixture from MidE1 should be approximately 20% in present-day samples from these groups.

Simulation Protocol: P1-P4

The focus of our study is on the impact of different demographic histories on P1-P4 as highlighted in Figure S1 (bottom). The global simulation scenario was thus adapted to specifically explore the impact of changing how P1 is related to P2.

For the Recent Bottleneck scenario, which closely follows the Marginalisation simulations of van Dorp et al, P1 splits from P2 relatively recently 20 generations before present, after which P1 undergoes an extreme instantaneous bottleneck that reduces the effective population size from 20,000 to 200 until present.

For the Ghost Admixture scenario, which largely follow the Remnants simulations of van Dorp et al, instead P1 splits from the ancestors of P2 1,700 generations in the past, maintaining an effective population size of 20,000. Additionally, P1 contributes migrants to P2 at a rate of 0.005 beginning at 300 generations ago and ending at 200 generations ago so that approximately 50% of P2 consists of migrants from P1 over this time period.

For the Recent Admixture scenario we generated a mixed population P2 as the result of an instantaneous admixture event occurring 15 generations ago between P1, P3 and P4. To do this we utilised the simulation technique employed by Price et al^10^ and also used in Leslie et al^11^. Using individuals simulated but not sampled from the Recent Bottleneck scenario we sample 20 individuals of P1, P3 and P4 in the proportions 50%, 35% and 15% to generate a haploid chromosome by sampling genetic distances in these proportions from an exponential of rate 0.15 (see Methods).

References

1.	Hellenthal, G. *et al.* A genetic atlas of human admixture history. *Science (80-. ).* **343,** 747–751 (2014).

2.	van Dorp, L. *et al.* Evidence for a Common Origin of Blacksmiths and Cultivators in the Ethiopian Ari within the Last 4500 Years: Lessons for Clustering-Based Inference. *PLOS Genet.* (2015). doi:10.1371/journal.pgen.1005397

3.	Marth, G. T., Czabarka, E., Murvai, J. & Sherry, S. T. The allele frequency spectrum in genome-wide human variation data reveals signals of differential demographic history in three large world populations. *Genetics* **166,** 351–372 (2004).

4.	Keinan, A., Mullikin, J. C., Patterson, N. & Reich, D. Measurement of the human allele frequency spectrum demonstrates greater genetic drift in East Asians than in Europeans. *Nat Genet* **39,** 1251–1255 (2007).

5.	Gutenkunst, R. N., Hernandez, R., Williamson, S. H. & Bustamante, C. D. Inferring the Joint Demographic History of Multiple Populations from Multidimensional SNP Frequency Data. *PLoS Genet* **5,** e1000695 (2009).

6.	Gronau, I., Hubisz, M. J., Gulko, B., Danko, C. G. & Siepel, A. Bayesian inference of ancient human demography from individual genome sequences. *Nat Genet* **43,** 1031–1034 (2011).

7.	Li, H. & Durbin, R. Inference of human population history from individual whole-genome sequences. *Nature* **475,** 493–496 (2011).

8.	Chen, G. K., Marjoram, P. & Wall, J. D. Fast and flexible simulation of DNA sequence data. *Genome Res.* **19,** 136–42 (2009).

9.	Pagani, L. *et al.* Ethiopian Genetic Diversity Reveals Linguistic Stratification and Complex Influences on the Ethiopian Gene Pool. *Am. J. Hum. Genet.* **91,** 83–96 (2012).

10.	Price, A. L. *et al.* Sensitive Detection of Chromosomal Segments of Distinct Ancestry in Admixed Populations. *PLoS Genet.* **5,** e1000519 (2009).

11.	Leslie, S. *et al.* The fine scale genetic structure of the British population. *Nature* **519,** 309–314 (2015).
